# Evo 2 Predicts Cardiomyopathy-Associated Variants and Elucidates Their Underlying Mechanisms

**DOI:** 10.64898/2026.05.15.725304

**Authors:** Atsumasa Kurozumi, Naoto Otsuka, Masamichi Ito, Toshinaru Kawakami, Takayuki Isagawa, Satoshi Kodera, Norihiko Takeda

## Abstract

**Background:** Although advances in next-generation sequencing have accelerated the identification of genetic variants in cardiomyopathy, interpreting variants of uncertain significance (VUS) remains a clinical challenge. Evo 2 is a high-resolution genomic artificial intelligence model capable of predicting pathogenicity across large sequence contexts and enabling mechanistic interpretation; however, its application in cardiovascular genetics is limited. Here, we evaluated the utility of Evo 2 for assessing the pathogenicity and underlying mechanisms of cardiomyopathy-associated variants.

**Methods:** We used Evo 2 to predict the pathogenicity of single-nucleotide variants in cardiomyopathy-related genes listed on ClinVar. We assessed the ability of the model to identify characteristic structural features in both coding and noncoding regions using internal representation such as embeddings, and to infer the molecular mechanisms of variants within these regions.

**Results:** Evo 2 demonstrated high predictive accuracy for pathogenicity, achieving an AUROC of 0.983 and an AUPRC of 0.915. Notably, sparse autoencoders (SAEs) from embeddings identified features corresponding to higher-order structural features, including coiled-coil and actin-binding domains characteristic of cardiomyopathy-related proteins, and accurately detected mutations known to disrupt these domains. The model recognized the binding motif of the cardiac-enriched transcription factor TBX5 with SAEs and accurately predicted a single-nucleotide polymorphism affecting TBX5 binding affinity after supervised fine-tuning.

**Conclusions:** Evo 2 demonstrated strong performance for both predicting pathogenicity and extracting biological features of cardiomyopathy-associated variants. It may represent a powerful emerging tool for evaluating VUS in cardiovascular medicine.

## 1. Introduction

Genetic testing for hypertrophic cardiomyopathy (HCM) and dilated cardiomyopathy (DCM) has become increasingly essential in clinical guidelines, informing accurate diagnosis (1,2), therapeutic decision-making, and prognostic stratification (3–5). Advances in next-generation sequencing have dramatically expanded the catalog of genetic variants (6,7); however, determining the clinical significance of rare or previously unreported variants remains challenging, often resulting in classification as variants of uncertain significance (VUS) (8). Although the American College of Medical Genetics and Genomics (ACMG) and Association for Molecular Pathology (AMP) guidelines strongly recommend functional validation using in vivo and in vitro assays (9), cardiovascular experiments are often time-and cost-intensive, particularly when generating induced pluripotent stem cell–derived cardiomyocytes (iPSCs).

To bridge this translational gap, artificial intelligence (AI) models have emerged as valuable tools for prioritizing variants before wet-lab validation and are recognized as supporting evidence in the ACMG/AMP guidelines (10). Genetic variants that may affect phenotypes include amino acid–altering variants in coding regions, regulatory variants in noncoding regions, and splice-altering variants located at the interface between coding and noncoding regions. Recent AI models have achieved high predictive performance for missense variants (e.g., AlphaMissense (11)) and noncoding variants (e.g., GPN-MSA (12)), alongside specialized approaches for predicting molecular mechanisms such as protein stability (11), splicing (13), and transcription factor (TF) binding motifs (14). Nevertheless, several limitations remain. Because it is difficult to analyze fine structural details while simultaneously considering the overall context, current approaches can predict global protein structural destabilization but still struggle to identify which specific secondary or tertiary amino acid structures are responsible for the instability. Moreover, accurate evaluation of noncoding regulatory elements, including enhancers, requires analysis of extensive long-range genomic context; however, transformer-based models often sacrifice single-nucleotide resolution to achieve broader sequence representation (15,16).

To address these challenges, the Arc Institute recently developed Evo (17) and Evo 2 (18) in 2024 and 2026, respectively. Using the StripedHyena2 architecture (19), these foundational models were trained on large-scale genomic sequences from diverse species, enabling the integration of long-range genomic context while preserving single-nucleotide resolution. This framework enabled two major advances. First, Evo 2 can predict pathogenicity by scoring variant effects in a zero-shot manner using likelihood-based scoring. For single-nucleotide variants in *BRCA1*, Evo 2 demonstrated predictive performance comparable to existing AI models for both coding and noncoding mutations. Second, Evo 2 offers interpretability through decomposition of model embeddings into sparse, high-dimensional representational representations using sparse autoencoders (SAEs) (20). This approach facilitates identification of biologically meaningful sequence features, including nucleotide patterns associated with protein structure and DNA motifs involved in molecular binding. Previous studies have identified features corresponding to protein secondary structures, such as α-helices and β-sheets, as well as binding motifs for multiple TFs. Furthermore, Evo 2 may enable detection of higher-order biological architectures, including tertiary protein structures and distal regulatory elements.

Importantly, organ-specific phenotypes are governed by tissue-specific regulatory networks. However, as Evo 2 was trained on diverse genomic sequences without tissue-specific annotations, its ability to predict cardiac-specific genetic variation and capture distinct cardiac structural features remains unclear.

Here, we investigated the utility of Evo 2 in cardiovascular genetics. Specifically, we evaluated whether Evo 2 can accurately predict the pathogenicity of variants in HCM- and DCM-associated genes and capture cardiac-specific genomic architectures, thereby providing insight into the molecular mechanisms underlying cardiomyopathy.

## 2. Results

We analyzed 4,457 pathogenic/likely pathogenic (P/LP) and 27,141 benign/likely benign (B/LB) SNVs across 24 ClinGen definitive HCM/DCM-associated genes from the ClinVar dataset. Evaluation of zero-shot Evo 2 likelihood scores for these SNVs (Figure 1) demonstrated strong predictive performance. Although CADD achieved the highest overall performance, Evo 2 outperformed all other models in both AUROC and AUPRC across all SNVs (Figure 2A). While positive predictive values (PPVs) varied among individual genes, overall performance remained consistent across the dataset (Supplementary Table 2). For coding mutations, Evo 2 maintained a high negative predictive value and outperformed several existing models, except for CADD (Figure 2B, Table 1). When analysis was restricted to non-synonymous mutations, Evo 2 achieved performance comparable to other models, although inferior to AlphaMissense (Figure 2C). Evo 2 also showed strong predictive performance for noncoding mutations, ranking second to CADD while remaining comparable to other approaches (Figure 2D, Table 1).

**Figure 1.**
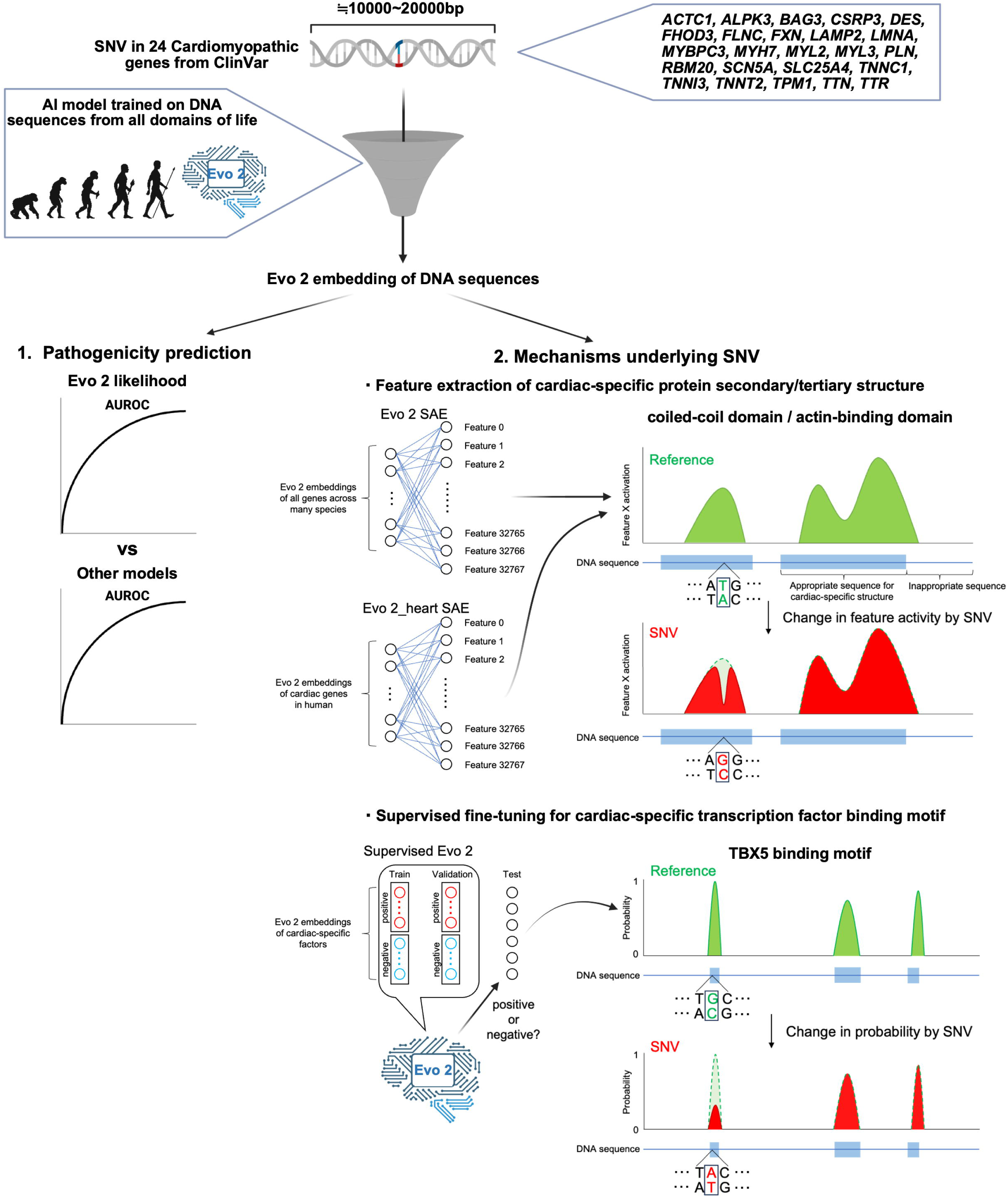
| Study design and workflow. In the first phase, SNVs in cardiomyopathy-associated genes were analyzed using Evo 2 to evaluate the predictive performance of likelihood-based pathogenicity scoring, with comparisons to other prediction models. In the second phase, to elucidate the underlying mechanisms of these SNVs, SAEs were applied to identify features capturing cardiac-specific elements, including higher-order protein structures and TF binding motifs. We then assessed whether SNVs affected the activation of these features. In parallel with the Evo 2 SAE, a cardiac-specific SAE (Evo 2_heart SAE) was developed using human heart–associated genes to enhance feature detection. Finally, we evaluated whether supervised fine-tuning of Evo 2 further improved predictive performance. AI, Artificial Intelligence; SAE, Sparse AutoEncoder; SNV, Single-Nucleotide Variant; TF, Transcription Factor

**Figure 2.**
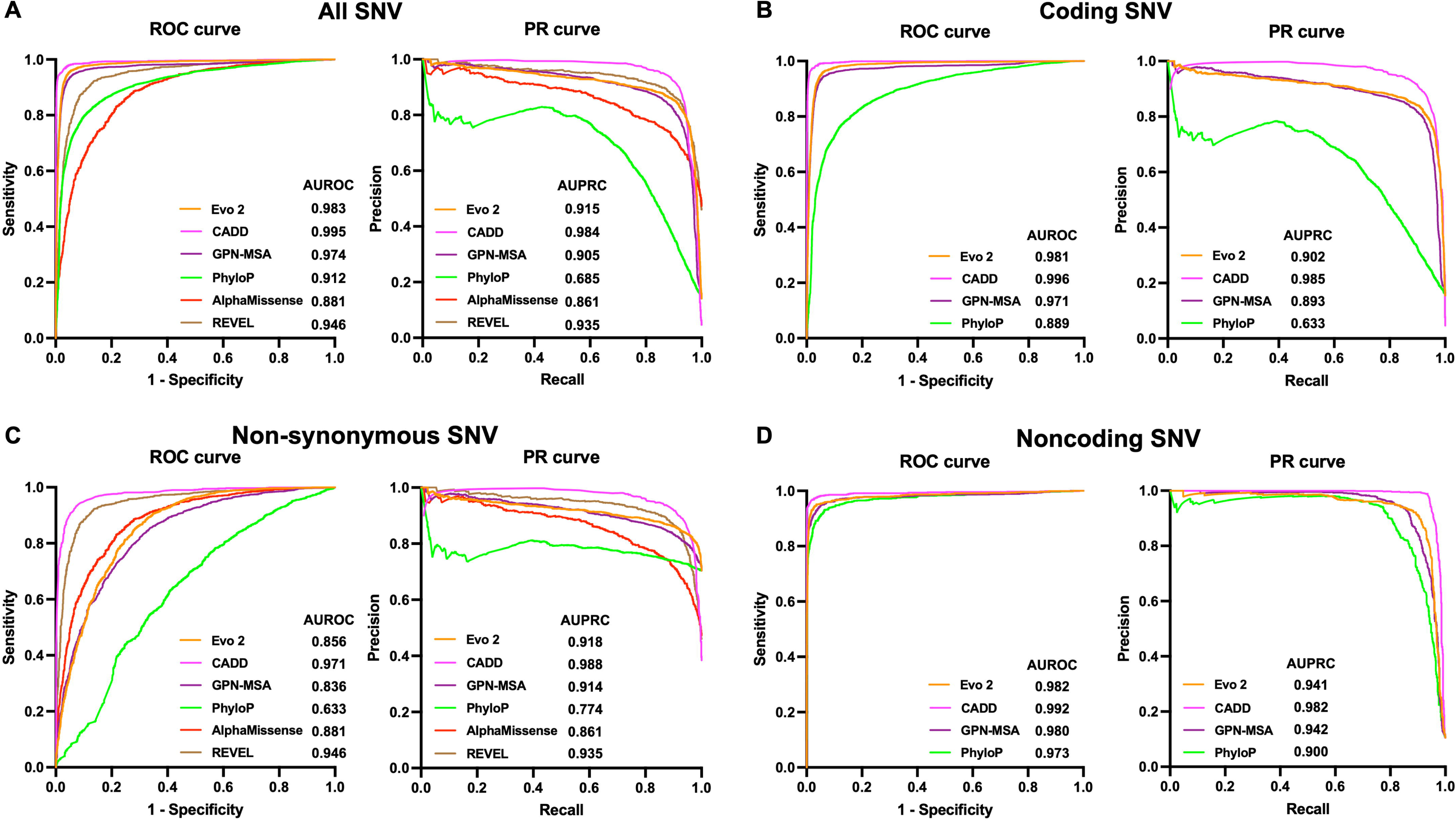
| Pathogenicity prediction of SNVs. A. ROC and PR curves for pathogenicity prediction of SNVs from ClinVar (n = 4,457 P/LP; 27,141 B/LB) across multiple prediction models (Evo 2, GPN-MSA, PhyloP, CADD, AlphaMissense, and REVEL). B. ROC and PR curves for coding SNVs (n = 3,535 P/LP; 19,305 B/LB). C. ROC and PR curves for non-synonymous SNVs (n = 3,521 P/LP; 1,478 B/LB). D. ROC and PR curves for noncoding SNVs (n = 922 P/LP; 7,836 B/LB). AI, Artificial Intelligence; B, Benign; LB, Likely Benign; LP, Likely Pathogenic; P, Pathogenic; PR, Precision–Recall; ROC, Receiver Operating Characteristic; SNV, Single-Nucleotide Variant

**Table 1.**
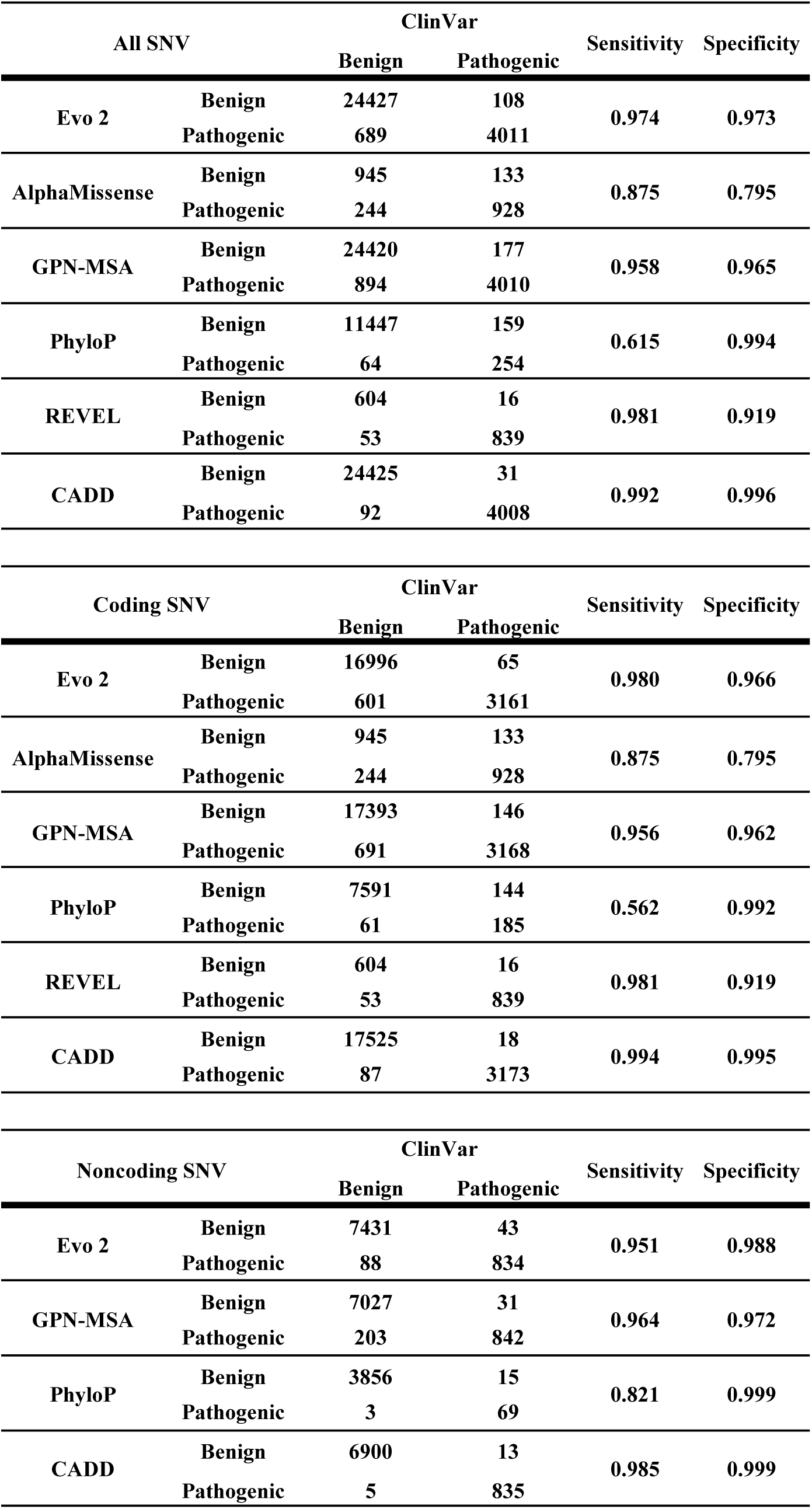
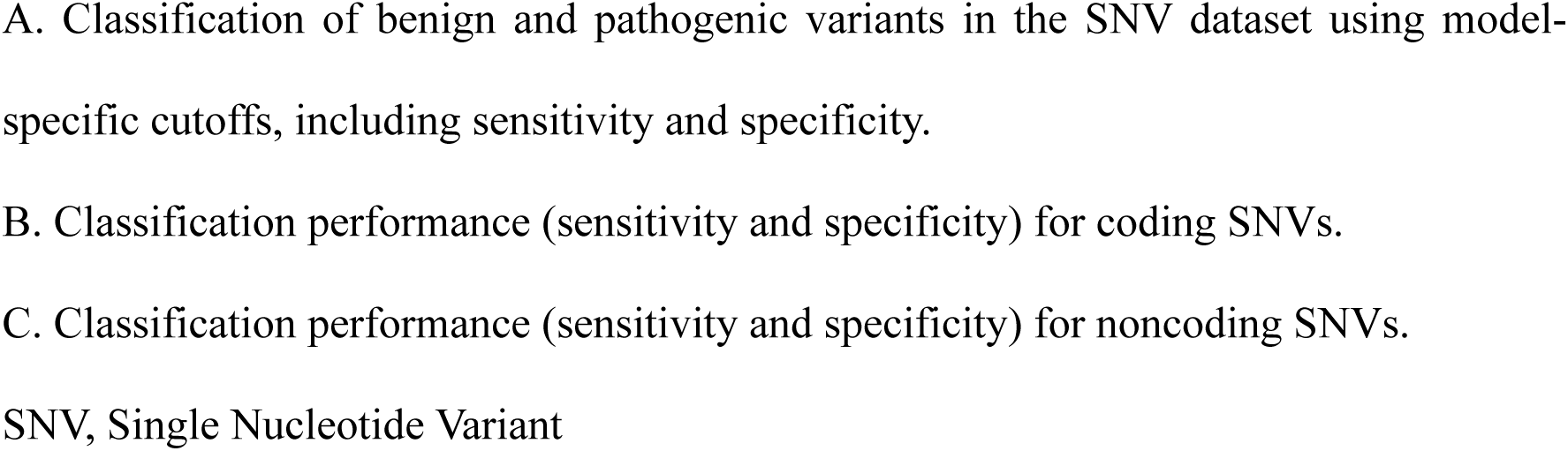
| Predictive performance for SNV pathogenicity.

| All SNV |  | ClinVar |  | Sensitivity | Specificity |
| --- | --- | --- | --- | --- | --- |
|  |  | Benign | Pathogenic |  |  |
| Evo 2 | Benign | 24427 | 108 | 0.974 | 0.973 |
|  | Pathogenic | 689 | 4011 |  |  |
| AlphaMissense | Benign | 945 | 133 | 0.875 | 0.795 |
|  | Pathogenic | 244 | 928 |  |  |
| GPN-MSA | Benign | 24420 | 177 | 0.958 | 0.965 |
|  | Pathogenic | 894 | 4010 |  |  |
| PhyloP | Benign | 11447 | 159 | 0.615 | 0.994 |
|  | Pathogenic | 64 | 254 |  |  |
| REVEL | Benign | 604 | 16 | 0.981 | 0.919 |
|  | Pathogenic | 53 | 839 |  |  |
| CADD | Benign | 24425 | 31 | 0.992 | 0.996 |
|  | Pathogenic | 92 | 4008 |  |  |
| Coding SNV |  | ClinVar |  | Sensitivity | Specificity |
|  |  | Benign | Pathogenic |  |  |
| Evo 2 | Benign | 16996 | 65 | 0.980 | 0.966 |
|  | Pathogenic | 601 | 3161 |  |  |
| AlphaMissense | Benign | 945 | 133 | 0.875 | 0.795 |
|  | Pathogenic | 244 | 928 |  |  |
| GPN-MSA | Benign | 17393 | 146 | 0.956 | 0.962 |
|  | Pathogenic | 691 | 3168 |  |  |
| PhyloP | Benign | 7591 | 144 | 0.562 | 0.992 |
|  | Pathogenic | 61 | 185 |  |  |
| REVEL | Benign | 604 | 16 | 0.981 | 0.919 |
|  | Pathogenic | 53 | 839 |  |  |
| CADD | Benign | 17525 | 18 | 0.994 | 0.995 |
|  | Pathogenic | 87 | 3173 |  |  |
| Noncoding SNV |  | ClinVar |  | Sensitivity | Specificity |
|  |  | Benign | Pathogenic |  |  |
| Evo 2 | Benign | 7431 | 43 | 0.951 | 0.988 |
|  | Pathogenic | 88 | 834 |  |  |
| GPN-MSA | Benign | 7027 | 31 | 0.964 | 0.972 |
|  | Pathogenic | 203 | 842 |  |  |
| PhyloP | Benign | 3856 | 15 | 0.821 | 0.999 |
|  | Pathogenic | 3 | 69 |  |  |
| CADD | Benign | 6900 | 13 | 0.985 | 0.999 |
|  | Pathogenic | 5 | 835 |  |  |

A. Classification of benign and pathogenic variants in the SNV dataset using model-specific cutoffs, including sensitivity and specificity.
B. Classification performance (sensitivity and specificity) for coding SNVs.
C. Classification performance (sensitivity and specificity) for noncoding SNVs.
SNV, Single Nucleotide Variant

To determine whether the performance of Evo 2 on coding mutations reflects an underlying representation of protein architecture, we applied sparse autoencoders (SAEs) to decode the model’s high-dimensional embeddings (Figure 1). We first focused on coiled-coil domains, key structural elements in cardiomyopathy-associated genes such as *LMNA* and *DES.* SAE analysis identified feature 20051 as strongly associated with coiled-coil domains (Figure 3A, Table 2, Supplementary Figure 1A). Non-synonymous mutations within coiled-coil domains produced significantly greater changes in feature 20051 activation than other mutations, suggesting that this feature may capture structural destabilization (Figure 3B). Consistent with this interpretation, the known destabilizing *LMNA* missense mutation c.644T>C (p.Leu215Pro) (21,22) caused a marked reduction in feature 20051 activation (Figure 3C).

**Figure 3.**
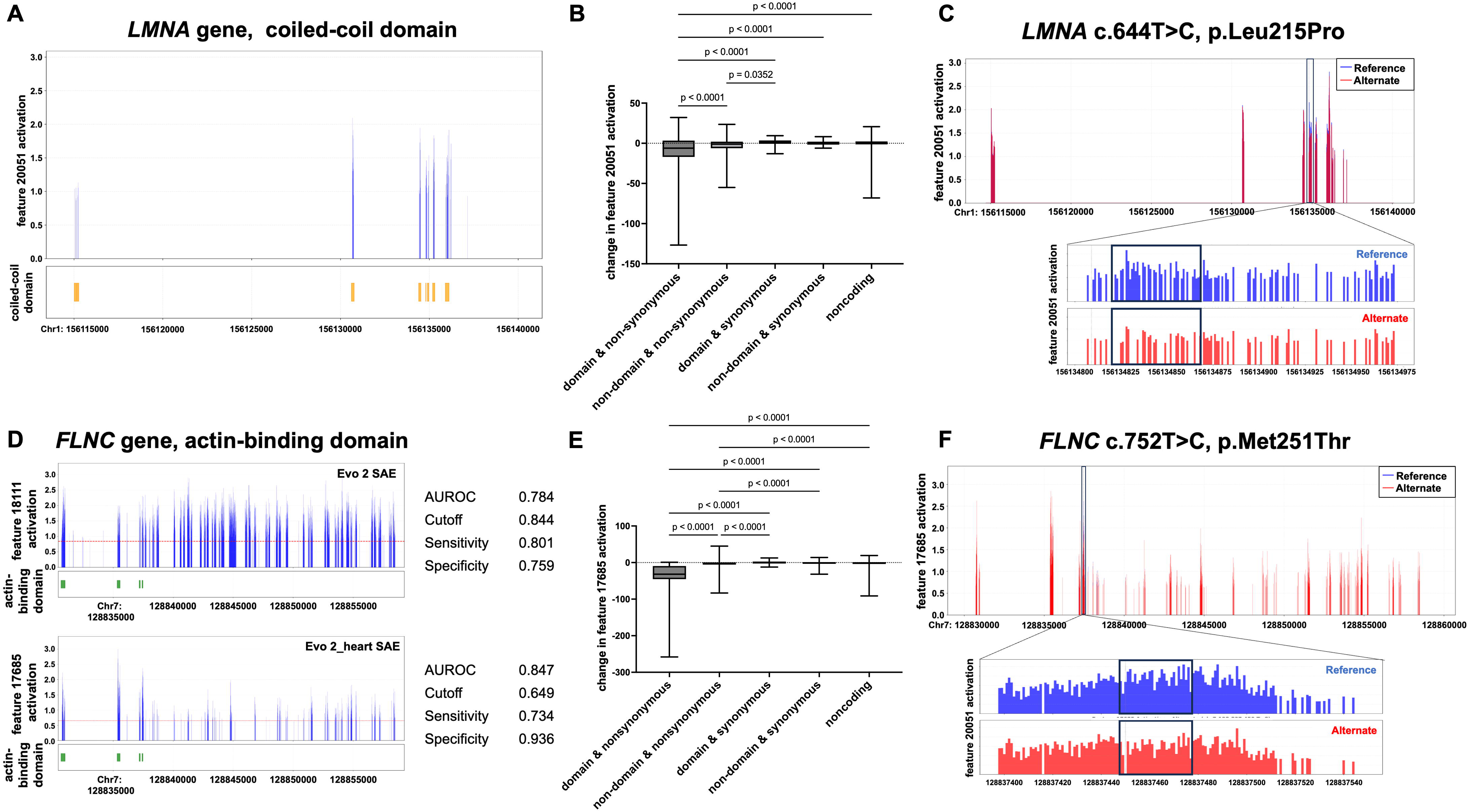
| Detection of characteristic exonic architectures and variant effect prediction. A. Activation of Evo 2 SAE feature 20051 (top) and corresponding coiled-coil domain regions (bottom) in *LMNA* genetic loci. B. Impact of SNVs in *LMNA* on Evo 2 SAE feature 20051 activation. Total activation changes across all *LMNA* bases were compared based on the types of mutations (non-synonymous and synonymous mutations, mutations within and outside the coiled-coil domain, as well as noncoding mutations). C. Changes in feature 20051 activation on the *LMNA* c.644T>C (p.Leu215Pro) variant. Blue, wild type sequence; Red, p.Leu215Pro mutant sequence. D. Activation of Evo 2 SAE feature 18111 and Evo 2_heart SAE feature 17685 in *FLNC* genetic loci. E. Impact of SNVs in *FLNC* on Evo 2_heart SAE feature 17685 activation. Total activation changes were compared based on the type of mutations (non-synonymous and synonymous mutations, mutations within and outside the actin-binding domain, as well as noncoding mutations). F. Changes in activation of Evo 2_heart SAE feature 17685 induced by the *FLNC* c.752T>C (p.Met251Thr) variant. Blue, wild tipe sequence; Red, p.Met251Thr mutant sequence. AUROC, Area Under the Receiver Operating Characteristic; *FLNC*, Filamin C; *LMNA*, Lamin A/C; SAE, Sparse AutoEncoder

**Table 2.**
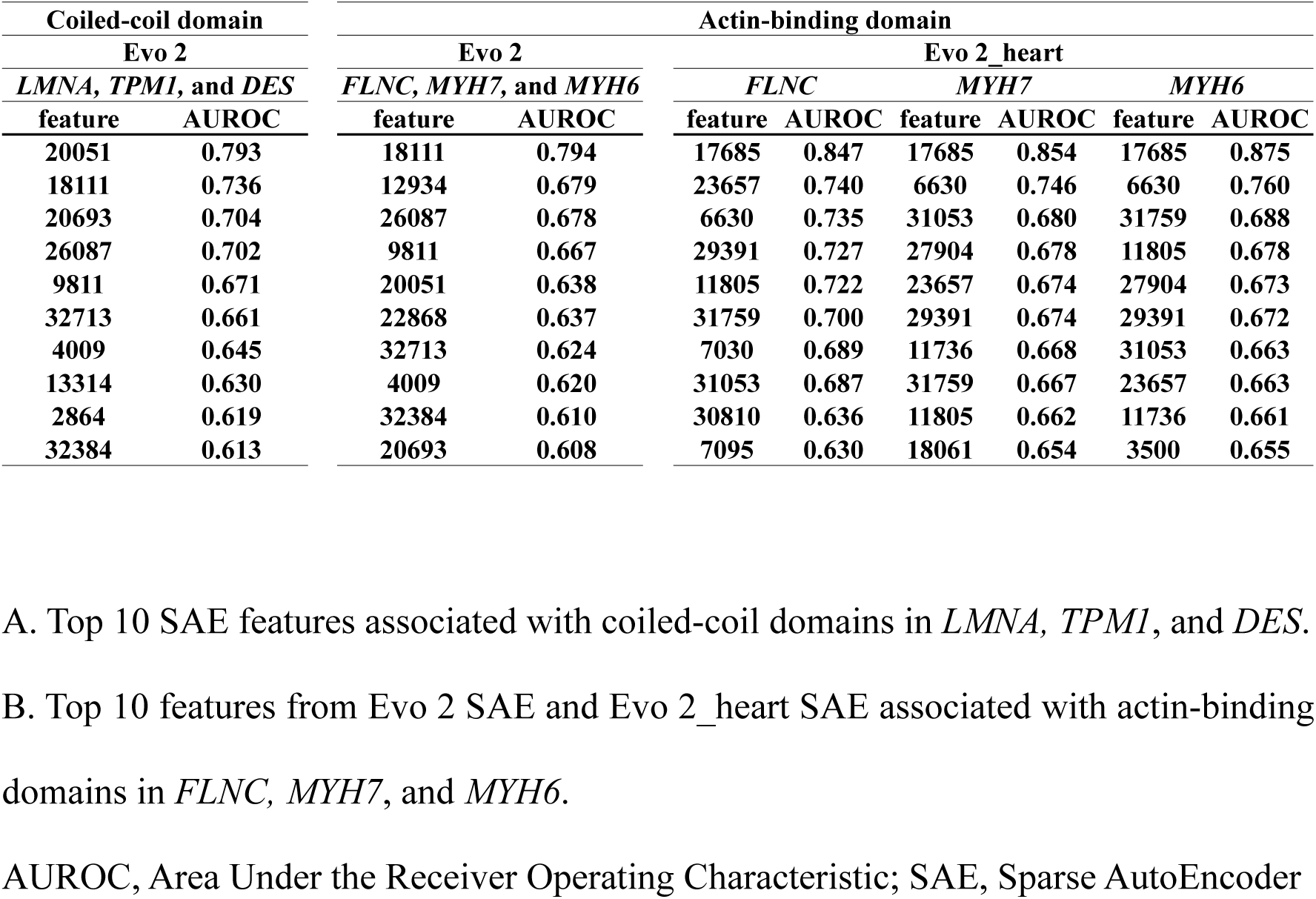
| Detection of heart-specific amino acid architectures.

In contrast, the original Evo 2 SAE showed limited ability to identify actin-binding domains, as the top-performing feature (feature 18111) demonstrated poor discriminative performance (Figure 3D). To address this limitation, we trained a cardiac-specific SAE (Evo 2_heart SAE) using 1,046 genes highly expressed in the human heart, identified from BioGPS (23). This organ-specific approach identified feature 17685, which substantially improved detection of actin-binding domain (Figure 3D, Table 2, Supplementary Figure 1B). Non-synonymous mutations within *FLNC* actin-binding domains induced significant change in feature 17685 activation (Figure 3E). Furthermore, the known destabilizing mutation *FLNC* c.752T>C (p.Met251Thr) (24) reduced the feature activation (Figure 3F). Together, these findings suggest that SAE feature activation patterns may help elucidate the molecular mechanisms underlying the pathogenicity of genetic variants.

Finally, we hypothesized that the ability of Evo 2 to predict the pathogenicity of noncoding mutations may reflect an implicit understanding of regulatory elements. We therefore focused on T-box transcription factor TBX5, a transcription factor known to regulate cardiomyopathy-associated genes including *MYH7* and *FLNC*. Variants within these loci can alter TBX5 binding affinity and contribute to cardiomyopathy (26). To investigate this possibility, we applied SAEs to predict TBX5 binding affinity from genomic sequences (Figure 1). Assuming that regions located within the target regions containing the consensus motif for TBX5 binding (TCACACCT) and the peak regions identified from previous ChIP-seq data for TBX5 in cardiomyocytes (ReMap TBX5 ChIP-seq data (25)) represent true TBX5 binding sites, we attempted to use SAE to identify regions that satisfy both of these conditions. Analysis of TBX5 ChIP-seq data revealed that binding peaks were enriched within ± 2,000 bp of transcription start sites (TSS) (Supplementary Figure 2B). Within these regions, we identified 49,978 TBX5 binding motifs (TBX5_motif_peak) and generated three additional datasets of equal size: TBX5_motif_nonpeak, TBX5_nonmotif_peak, and TBX5_nonmotif_nonpeak.

SAE analysis identified feature 9152, which exhibited significantly higher activation in TBX5_motif_peak sequences (Figure 4A, Supplementary Table 3). Although this feature showed poor discrimination for the TBX5_nonmotif_peak dataset, its activation patterns were comparable between the TBX5_motif_nonpeak and TBX5_nonmotif_nonpeak datasets (Figure 4A). These findings suggest that feature 9152 is not activated solely by the canonical 8-bp TBX5 binding motif, but also captures the surrounding genomic context. Evo 2_heart SAE further improved detection of TBX5-associated regulatory features (Supplementary Table 3).

**Figure 4.**
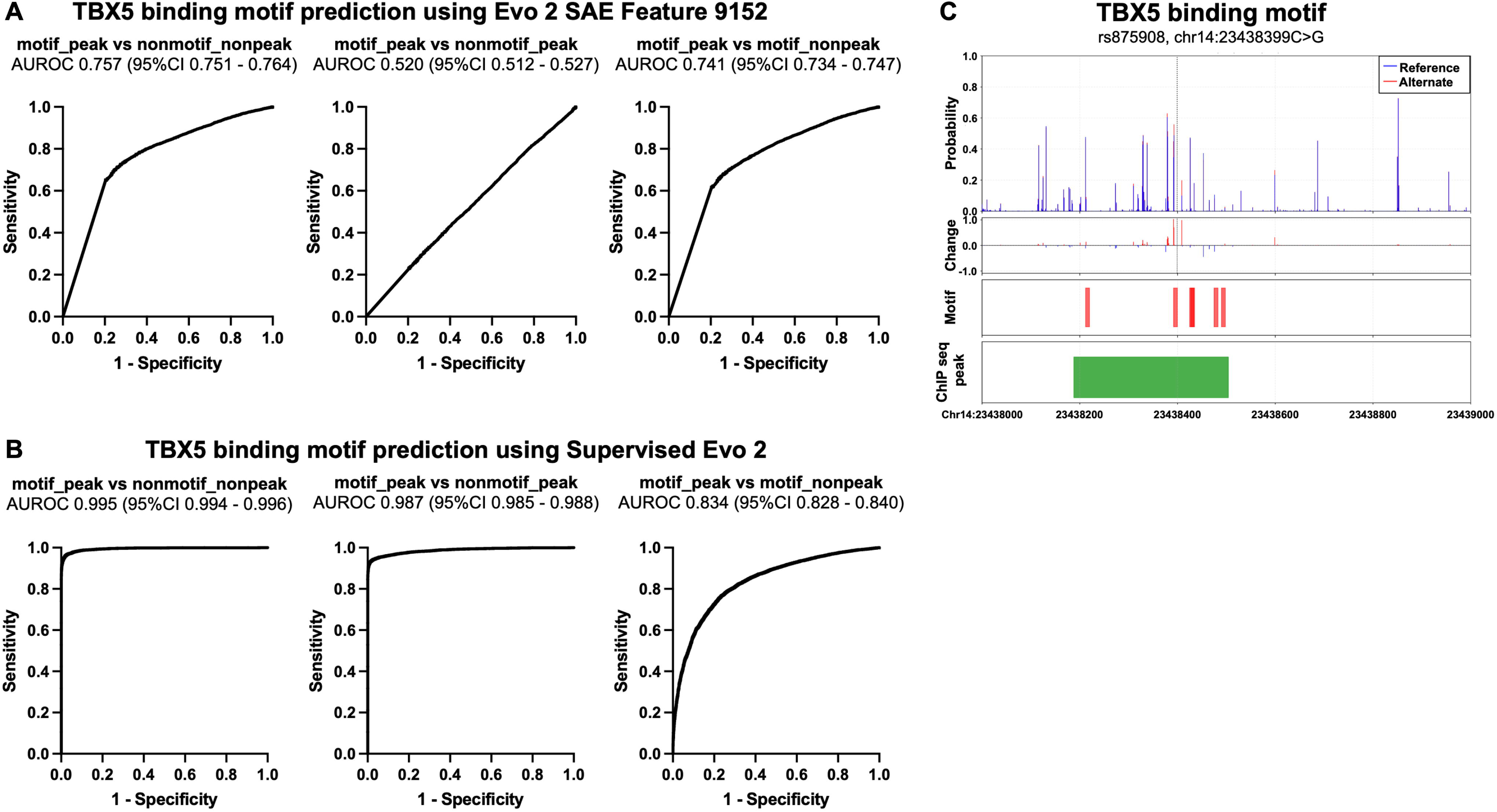
| Detection of functional noncoding sequences and variant effect prediction. A. ROC curves evaluating the predictive performance of Evo 2 SAE feature 9152 activation distinguishing TBX5_motif_peak sequences from three other sequence groups (TBX5_nonmotif_nonpeak, TBX5_nonmotif_peak, and TBX5_motif_nonpeak). B. ROC curves evaluating the predictive performance of the supervised Evo 2 model in distinguishing TBX5_motif_peak sequences from the other sequence groups. C. Changes in activation of supervised Evo 2 TBX5 binding probability induced by a SNP in *MYH7* promoter region (rs875908). Blue, wild type sequence; Red, mutant sequence. AUROC, Area Under the Receiver Operating Characteristic; ROC, Receiver Operating Characteristic; SAE, Sparse AutoEncoder

We next developed a supervised Evo 2 classifier for TBX5 binding prediction using embeddings of the four datasets described above. The model achieved AUROC values of 0.995, 0.987, and 0.834 on the TBX5_nonmotif_nonpeak, TBX5_nonmotif_peak, and TBX5_motif_nonpeak datasets, respectively (Figure 4B), while maintaining predictive performance at greater distances from the TSS (Supplementary Figure 2C). The supervised Evo 2 TF-binding model also successfully predicted altered binding affinity for rs875908 (26), a known single-nucleotide polymorphism located 3,000 bp upstream of the *MYH7* TSS (Figure 4C).

## 3. Discussion

In this study, we demonstrated that Evo 2 effectively predicts the pathogenicity of SNVs in cardiomyopathy-associated genes while also enabling the detection of abnormalities in cardiac-specific amino acid structures and TF-binding motifs.

Evo 2 utilizes the convolutional multi-hybrid architecture StripedHyena2 (19), which combines multiple types of operators to improve the efficiency of both training and inference. It was trained on a total of 8.8 trillion base pairs across 40 billion sequences, each up to one million base pairs in length, encompassing both prokaryotic and eukaryotic DNA sequences. As an autoregressive language model trained to predict the next base in a sequence, Evo 2 can model long-range dependencies while preserving single-nucleotide resolution. Evo 2 learns sequence likelihood landscapes across diverse DNA sequences and can predict the effects of genetic variants on biological function in a zero-shot manner without task-specific fine-tuning or supervision. Leveraging its likelihood estimation capability, variant effects can be quantified by changes in log-likelihood (Δlikelihood) between variant and reference sequences, enabling pathogenicity prediction.

Our zero-shot analysis demonstrated strong performance for both coding and noncoding SNVs. Considering that CADD, AlphaMissense, and REVEL are supervised models, the zero-shot performance of Evo 2 is particularly noteworthy. PPV were lower for specific genes (*FHOD3, MYL3,* and *RBM20*). However, according to LOEUF, sequence constraint in these genes was not lower than that in other genes. These genes tended to harbor more benign than pathogenic missense SNVs, suggesting that genes predominantly comprising benign variants may be more susceptible to false-positive predictions. Nevertheless, because sensitivity and negative predictive value remained high even for these genes, the model may still be useful as a screening tool.

In the cardiovascular field, in vitro validation often requires induced pluripotent stem cell–derived cardiomyocytes, which are associated with substantial cost and labor, thereby making validation of genetic variants challenging. Consequently, highly accurate and interpretable in silico models are critical prior to wet-lab validation. Notably, Evo 2, combined with analyses of internal representation such as SAEs, successfully detected cardiac-specific structures, highlighting its potential to elucidate variant mechanisms in silico. We successfully detected coiled-coil domains—supersecondary α-helical structures essential for mechanical resilience in the heart (27)—and identified destabilizing mutations within these domains. However, Evo 2 SAEs, which were limited to approximately 30,000 features, showed limited ability to detect more complex, tissue-specific architectures such as actin-binding domains. A cardiac-specific SAE trained on heart-enriched genes overcame this limitation, enabling the identification of features corresponding to these structures. Because Evo 2 was trained on DNA sequences from a wide range of species without restriction to specific organs, our results suggest that this approach may enable identification of diverse organ-specific structural features, even for complex protein architectures. It is well established that phenotypic outcomes vary depending on the position of amino acid substitutions (28,29); therefore, because Evo 2 captures amino acid–related features, it may ultimately help predict the contribution of genetic variants to phenotypes such as heart failure and arrhythmia.

Transcription factor binding remains incompletely understood. While zero-shot Evo 2 captured motif patterns, supervised fine-tuning enabled highly accurate prediction of TBX5-binding motifs. Intriguingly, we identified a feature capable of predicting TBX5 binding based on flanking nucleotide sequences, even when the consensus motif itself was identical. It has become increasingly recognized that sequences outside core TF binding motifs also play important roles (30), suggesting the potential for further elucidation of TF-binding mechanisms.

In addition, although the performance of PromoterAI declines with increasing distance from the TSS (31), Evo 2 showed no substantial decline in performance. This likely reflects its strength in capturing long-range sequence dependencies. Furthermore, although TF binding is strongly influenced by epigenomic factors such as chromatin accessibility, histone modifications, and interactions with other proteins (32), our results suggest that TF binding may be partially predictable from genomic sequence information alone.

In this study, we employed both SAEs and supervised fine-tuning to identify cardiac-specific features. Although supervised fine-tuning was effective for predicting TBX5-binding motifs, SAEs were also indispensable. Supervised learning depends on labeled data and is therefore well suited for predicting known structures; however, SAEs are essential for capturing features in sequences such as DNA, where many functional elements remain uncharacterized. When the mechanism by which genetic variants contribute to phenotypes are unknown, identifying features to exhibit large changes and linking them to variants with known mechanisms may help elucidate the underlying mechanism.

Several limitations should be noted. First, we relied on ClinVar annotations as labels. Second, our analysis was restricted to SNVs; although Evo 2 can compute likelihoods for insertions and deletions, applying Δlikelihood thresholds to such variants requires caution. Third, assuming that all non-synonymous mutations within coiled-coil or actin-binding domains cause structural instability may lead to overestimation. Finally, although we defined TBX5-binding sites as motif-containing peak regions, previous studies have shown that TBX5 can bind outside canonical motifs through indirect binding mediated by co-binding with other TFs (33,34). Therefore, the training data may not fully represent authentic TBX5-binding sites.

In conclusion, Evo 2 accurately predicts the pathogenicity of cardiomyopathy-associated variants and captures cardiac-specific structural features. Moreover, based on the performance of Evo 2 in predicting the effects of variants in noncoding regions and TF binding to DNA sequences, our results suggest that TF binding in the heart may be partially elucidated using genomic sequence information alone. Evo 2 may serve as a powerful in silico tool for evaluating VUS and elucidating their underlying mechanisms prior to in vitro and in vivo validation.

## Author contributions

A.K., M.I., and S.K. conceived and designed the study. A.K. and N.O. collected and curated the datasets, performed bioinformatic and statistical analyses, and developed the AI-based analytical framework. A.K. interpreted the results and drafted the manuscript. All authors contributed to data interpretation, critically revised the manuscript for important intellectual content, and approved the final version of the manuscript.

## Acknowledgements

None.

## 5. Methods

### SNV Dataset

We curated 24 definitive genes associated with hypertrophic cardiomyopathy (HCM) and dilated cardiomyopathy (DCM) (*ACTC1, ALPK3, BAG3, CSRP3, DES, FHOD3, FLNC, FXN, LAMP2, LMNA, MYBPC3, MYH7, MYL2, MYL3, PLN, RBM20, SCN5A, SLC25A4, TNNC1, TNNI3, TNNT2, TPM1, TTN,* and *TTR*) based on the Clinical Genome Resource (ClinGen; accessed August 2025). From these genes, we extracted single-nucleotide variants (SNVs) annotated as Pathogenic (P), Likely Pathogenic (LP), Benign (B), or Likely Benign (LB) from the ClinVar database (NCBI; accessed August 2025).

### Scoring of Evo 2 and other Models

For zero-shot scoring with Evo 2, we used an 8,192-bp context window centered on the mutated nucleotide. The likelihoods of both the reference and mutant sequences were calculated as the product of predicted probabilities for all tokens within the window. The variant score was defined as the difference in log-likelihoods:

Δlikelihood = log P(mutant sequence) - log P(reference sequence)

We also computed SNV scores using additional models, including GPN-MSA (12), PhyloP (35), CADD (36), AlphaMissense (11), and REVEL (37). Model performance was evaluated using receiver operating characteristic (ROC) curves and precision–recall (PR) curves, with P/LP variants assigned as positive (label 1) and B/LB variants as negative (label 0). Cutoff values for pathogenic, ambiguous, and benign classifications were determined at recall > 0.9 for Evo 2, GPN-MSA, CADD, and AlphaMissense (Supplementary Table 1). ClinGen-recommended moderate thresholds were applied for PhyloP and REVEL.

### Coiled-coil and Actin-binding Domain Analysis

Amino acid sequences corresponding to the coiled-coil domains of *TPM1, LMNA,* and *DES,* and the actin-binding domains of *FLNC, MYH7*, and *MYH6*, were obtained from UniProt (38) and mapped to their genomic coordinates (GRCh38). Embeddings were extracted using an 8,192-bp window centered on the midpoint of each exon or intron, and feature activations were computed using the SAE. Exons and introns longer than 8,192 bp were segmented prior to analysis.

To identify salient SAE features, AUROC values were calculated by comparing feature activations between domain regions (label 1) and non-domain regions (label 0). Activations of the selected features were compared across coiled-coil and actin-binding domains, as well as randomly selected exons from other cardiomyopathy-associated genes.

Changes in feature activation induced by SNVs were also computed and visualized. ClinVar SNVs (P/LP and B/LB) in *LMNA* and *FLNC* were stratified according to genomic location (domain vs. non-domain) and mutation type (synonymous, non-synonymous, and noncoding), and corresponding feature activations were evaluated.

### TF Binding Motif Analysis by SAEs and Supervised Classifications

To identify salient SAE features associated with TBX5 binding motifs, we compared mean 8-bp feature activations between the TBX5_motif_peak (label 1) and TBX5_nonmotif_nonpeak (label 0) datasets and selected features with the highest AUROC values. These features were subsequently evaluated across the remaining two datasets.

Furthermore, we developed a supervised Evo 2–based classifier to predict TBX5 binding motifs. For each 8-bp sequence across all datasets, embeddings were extracted from layer 26 (blocks.26.mlp.l3) using an 8,192-bp context window. Forward and reverse-complement embeddings (4,096 dimensions each) were concatenated to generate a 65,536-dimensional feature vector. The classifier consisted of a five-layer feedforward neural network with the following architecture:

Input (65,536) → Linear (8192)→ ReLU→ BatchNorm→ Dropout (0.3)→ Linear (2048)→ ReLU→BatchNorm→Dropout (0.3)→Linear (512)→ReLU→BatchNorm→Dropout (0.3)→ Linear(128)→ReLU→BatchNorm→Dropout (0.15)→Linear (1)

The classifier was trained to distinguish TBX5_motif_peak (label 1) from all other datasets (label 0) using BCEWithLogitsLoss with class weighting. Optimization was performed using the Adam optimizer (learning rate: 1 × 10³, weight decay: 1 × 10, batch size: 256). Training stability was ensured via early stopping (patience: 7 epochs) and learning rate decay (factor: 0.5, patience: 3 epochs) based on validation performance.

