## Supplementary figures and images for "Evo 2 Predicts Cardiomyopathy-Associated Variants and Elucidates Their Underlying Mechanisms"

### Supplementary Figure 1

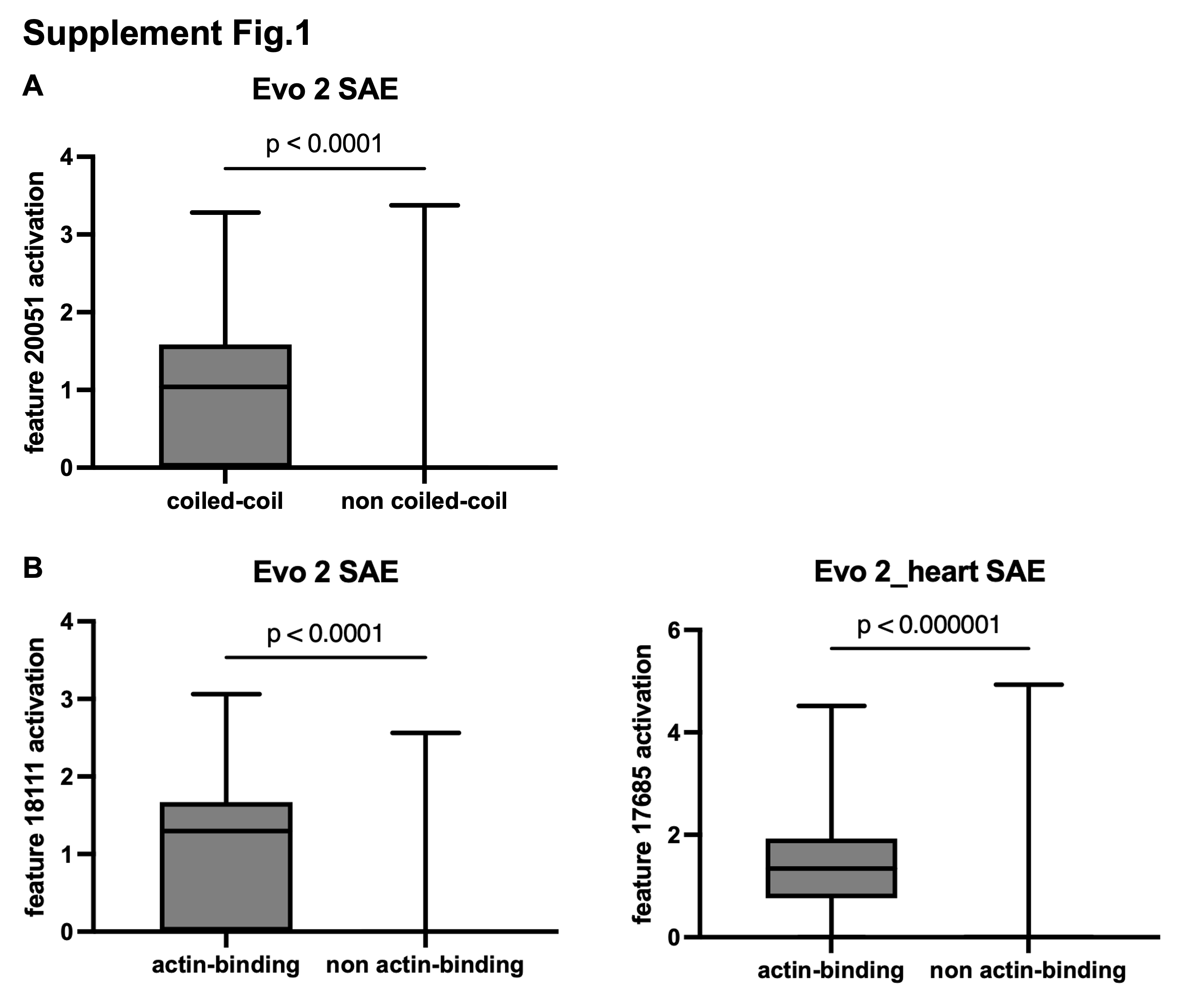

### Supplementary Figure 2

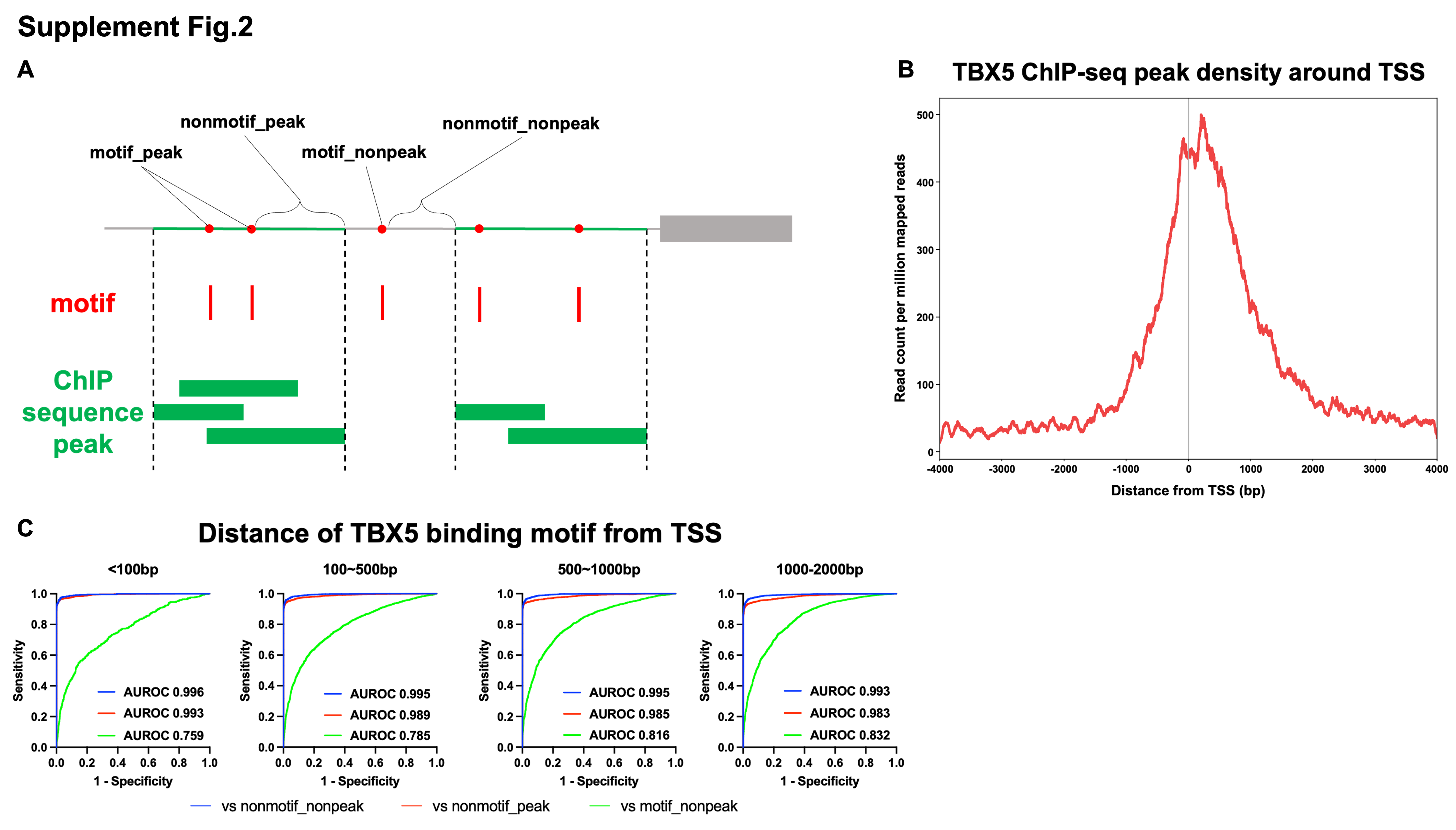
