## Supplementary Figure Legend for "Evo 2 Predicts Cardiomyopathy-Associated Variants and Elucidates Their Underlying Mechanisms"

**Supplementary Figure Legends**

**Supplementary Figure 1 | Prediction of heart-associated amino acid structures.**

A. Comparison of Evo 2 SAE feature 20051 activation between coiled-coil domains (in *LMNA, TPM1,* and *DES*) and exonic regions lacking coiled-coil domains (in other cardiomyopathy-associated genes).

B. Comparison of Evo 2_heart SAE feature 17685 activation between actin-binding domains (in *FLNC, MYH7,* and *MYH6*) and exonic regions lacking actin-binding domains (in other cardiomyopathy-associated genes).

SAE, Sparse AutoEncoder

**Supplementary Figure 2 | TF ChIP-seq peak distribution and binding motif prediction.**

A. Schematic representation and definition of regions related to TF binding motifs and ChIP-seq peaks. Red dots indicate consensus binding motifs of TF and green lines indicate ChIP-seq peaks on genomic sequences. Genomic resions are categorized into four types: TBX5_motif_peak, TBX5_nonmotif_peak, TBX5_motif_nonpeak, and TBX5_nonmotif_nonpeak.

TBX5_motif_peak, sequences with TBX5 consensus motif (TCACACCT) and within the peak regions previously identified by TBX5 ChIP-seq experiments using cardiomyocytes.

TBX5_nonmotif_peak, sequences without TBX5 consensus motif (TCACACCT) and within of the peak regions previously identified by TBX5 ChIP-seq experiments using cardiomyocytes.

TBX5_motif_nonpeak, sequences with TBX5 consensus motif (TCACACCT) and outside of the peak regions previously identified by TBX5 ChIP-seq experiments using cardiomyocytes.

TBX5_nonmotif_nonpeak, sequences without TBX5 consensus motif (TCACACCT) and within the peak regions previously identified by TBX5 ChIP-seq experiments using cardiomyocytes.

B. Distribution of TBX5 ChIP-seq peaks relative to TSS. Peak region information was extracted from the ReMap database.

C. ROC curves evaluating the predictive performance of the supervised Evo 2 model in distinguishing TBX5_motif_peak sequences from the other sequence groups (TBX5_nonmotif_nonpeak, TBX5_nonmotif_peak, and TBX5_motif_nonpeak) across distance bins from the TSS (0 – 100 bp, 100 – 500 bp, 500 – 1,000 bp, and 1,000 – 2,000 bp). AUROC, Area Under the Receiver Operating Characteristic curve; ChIP, Chromatin Immunoprecipitation; SAE, Sparse AutoEncoder; TF, Transcription Factor; TSS, Transcription Start Site.
