## Supplementary Method for "Evo 2 Predicts Cardiomyopathy-Associated Variants and Elucidates Their Underlying Mechanisms"

Supplementary Methods

Evo2 SAE

To compute SAE activations, we used a publicly available sparse autoencoder from Goodfire (sae-layer26-mixed-expansion_8-k_64.pt), trained on embeddings from the mixed module at layer 26 of Evo 2. The SAE comprises 32,768 latent dimensions (expansion factor 8, k = 64). Within an 8,192-bp context window centered on each base, embedding vectors for each nucleotide position were extracted from layer 26 of Evo 2 and input into the SAE to compute feature activations across all 32,768 dimensions at each position.

Evo2_heart SAE

Using BioGPS (1), we identified genes with cardiac expression levels greater than threefold higher than in other tissues. For these heart-enriched genes, embeddings were extracted from Evo 2 block 26 (blocks.26.mlp.l3) using an 8,192-bp window centered at each position across exonic and intronic regions. We developed a cardiac-specific SAE (Evo2_heart SAE; https://huggingface.co/harari/tbx5-sae-k128) using the BatchTopKTiedSAE architecture. The model was trained with mean squared error (MSE) loss, with an input dimension of 4,096, a hidden dimension of 32,768 (expansion factor 8), and k = 128.

Datasets of TF Binding Motif Analysis

TBX5 chromatin immunoprecipitation sequencing (ChIP-seq) data (GSE81585, GSE85628) were obtained from ReMap (2). Based on the Position Weight Matrix (PWM) of TBX5 from JASPAR, sequences with p < 0.005 relative to genome-wide background frequencies were identified using HOMER (3) as TBX5-binding motifs. We restricted the analysis to sequences within ±2,000 bp of transcription start sites (TSS), based on their distribution. Motif-containing sequences within ChIP peaks were defined as the TBX5_motif_peak dataset. Motif sequences located outside ChIP peaks were defined as the TBX5_motif_nonpeak dataset and were randomly sampled. Additionally, random 8-bp sequences with p > 0.005 within ±2,000 bp of TSS were sampled and categorized as TBX5_nonmotif_peak (within ChIP peaks) or TBX5_nonmotif_nonpeak (outside peaks) (Supplementary Figure 2A). All four datasets were constructed to have equal sample sizes.
