## Supplementary Table for "Evo 2 Predicts Cardiomyopathy-Associated Variants and Elucidates Their Underlying Mechanisms"

**Supplementary Table 1 | Cutoff values for computational models.**

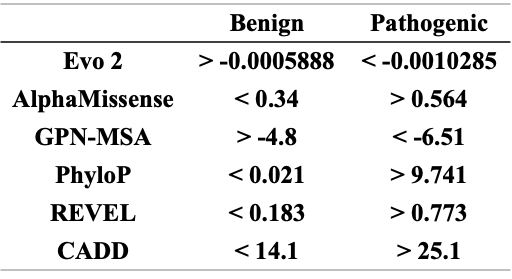

Pathogenic and benign classification cutoffs for Evo 2, GPN-MSA, CADD, and AlphaMissense were defined at recall > 0.9. Cutoffs for PhyloP and REVEL were based on ClinGen moderate recommendation thresholds.

**Supplementary Table 2 | Gene-level pathogenicity prediction.**

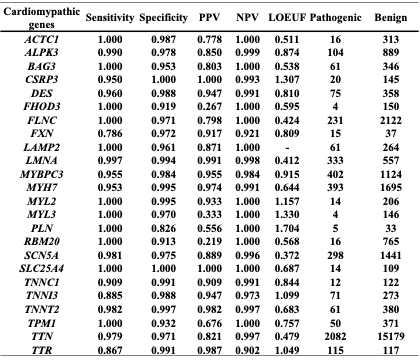

Sensitivity, specificity, PPV, and NPV of zero-shot Evo 2 predictions for SNVs in each cardiomyopathy-associated gene, along with LOEUF scores and variant classifications.

LOEUF, Loss of function Observed/Expected Upper bound Fraction; NPV, Negative Predictive Value; PPV, Positive Predictive Value; SNV, Single Nucleotide Variant.

**Supplementary Table 3 | Prediction of TBX5 binding motifs with SAE.**

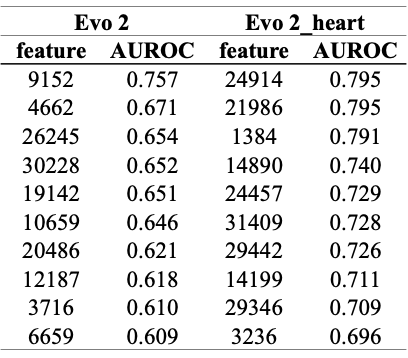

Top 10 features from Evo 2 SAE and Evo 2_heart SAE distinguishing TBX5_motif_peak from TBX5_nonmotif_nonpeak sequences.

AUROC, Area Under the Receiver Operating Characteristic; SAE, Sparse AutoEncoder.
